# 11 Million SNP Reference Profiles for Identity Searching Across Ethnicities, Kinship, and Admixture

**DOI:** 10.1101/321190

**Authors:** Brian S. Helfer, Darrell O. Ricke

## Abstract

High throughput sequencing (HTS) of single nucleotide polymorphisms (SNPs) provides additional applications for DNA forensics including identification, mixture analysis, kinship prediction, and biogeographic ancestry prediction. Public repositories of human genetic data are being rapidly generated and released, but the majorities of these samples are de-identified to protect privacy, and have little or no individual metadata such as appearance (photos), ethnicity, relatives, etc. A reference in silico dataset has been generated to enable development and testing of new DNA forensics algorithms. This dataset provides 11 million SNP profiles for individuals with defined ethnicities and family relationships spanning eight generations with admixture for a panel with 39,108 SNPs.

## Introduction

The current FBI National DNA Index System (NDIS) database has over 16 million profiles^1^. Illumina recently introduced the ForenSeq^2^ sequencing panel that includes single nucleotide polymorphisms (SNPs) in addition to short tandem repeats (STRs). This expanded panel of STR and SNP profiles enables biogeographic ancestry (BGA) and phenotype (externally visible traits – EVTs) predictions. Others have pioneered using SNPs for BGA prediction^3,4^, EVTs^5^, mixture analysis^6,7^, and more. Future HTS/massively parallel sequencing (MPS) panels will scale to thousands and tens of thousands of loci selected for multiple forensics applications^8^.

There is currently an unmet need for a reference dataset providing SNP information across different ethnic groups with defined relationships for millions of individuals. There are multiple public repositories [Alfred, dbSNP, 1,000 genomes, etc.] of human genetic variants^9–11^, but none of them provide sufficient data yet to meet the needs for developing some DNA forensics algorithms. To model a SNP dataset on the scale of NDIS, 11 million in silico profiles were generated using well-characterized SNPs; ethnicities and kinship information were tracked across generations. From the initial starting four ethnic founding populations, eight generations of descendants were generated to create realistic population data including multiple marriages with half-siblings, admixture, and children of consanguineous marriages represented. An additional set of unrelated individuals from these four ethnicities was also generated for comparison. This dataset provides great utility for testing identity searching, kinship prediction, and biogeographic ancestry predication methods. The incorporated SNPs were selected from a set of loci that were well characterized across multiple ethnicities.

## Method

### Database Generation

SNPs were selected from the Allele Frequency Database (ALFRED)^12^, which contains large numbers of SNPs that are well-characterized across ethnicities; a total of 39,108 SNPs were characterized across African Americans, Estonians, Koreans, and Palestinians. This information was used to create four initial in silico populations of pure ancestry, each containing 15,000 individuals. When multiple sources were listed for a single locus, the median minor allele frequency was assigned to the SNP. The gender of each individual was assigned such that 
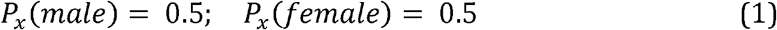
the probability that individual x is male is 0.5; and, the probability that individual x is female is also 0.5. Minor allele information was assigned using the allele frequencies provided by ALFRED. Eight more generations were created using these four ethnic populations. A set of unrelated individuals were also generated with the same four initial ethnic groups using a different seeding of the random number generator. This results in a starting generation of individuals that will have relatives and an independent set of individuals with no relatives.

Within each subsequent generation a constrained pairing of a male and female from the previous generation produced offspring. The pairings generated between individuals were constrained based on previous U.S. census information^13^. Within each generation, the probability that an individual would marry and how many times, given they are male is:
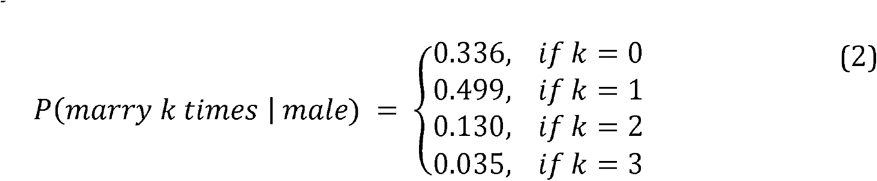
and the probability that an individual would marry, given they are female is:
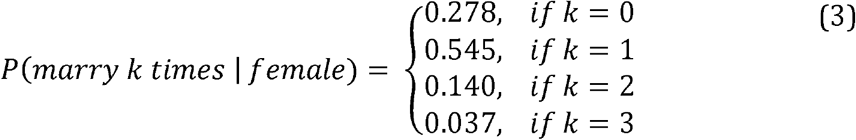

This information provides the probability that a person would get married, zero, one, two, or three or more times.

Furthermore, an ethnic intermarriage rate 
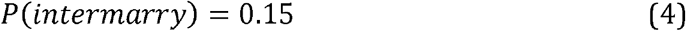
was set based on a poll conducted by Pew Research Center on social trends^14^. If an individual were the product of intermarriage, they would be given a uniform probability of marrying all individuals of the opposite gender, regardless of their ethnicity. All couples that married produced four offspring. Each child was created independently with their two alleles drawn randomly from their parents at each locus. This procedure was followed for the remaining eight generations. No modeling of allele phasing was included. A total of 11,164,912 profiles were generated.

## Results

One application of this dataset is for testing and validation of identity search algorithms. The SNP comparison times (not including dataset reading time) for the MIT Lincoln Laboratory FastID^15^ tool using a single computational thread on a panel of 5,000 SNPs is shown in Figure 1. The FastID algorithm leverages this in silico dataset as test data for validation and timing.

**Figure 1.**
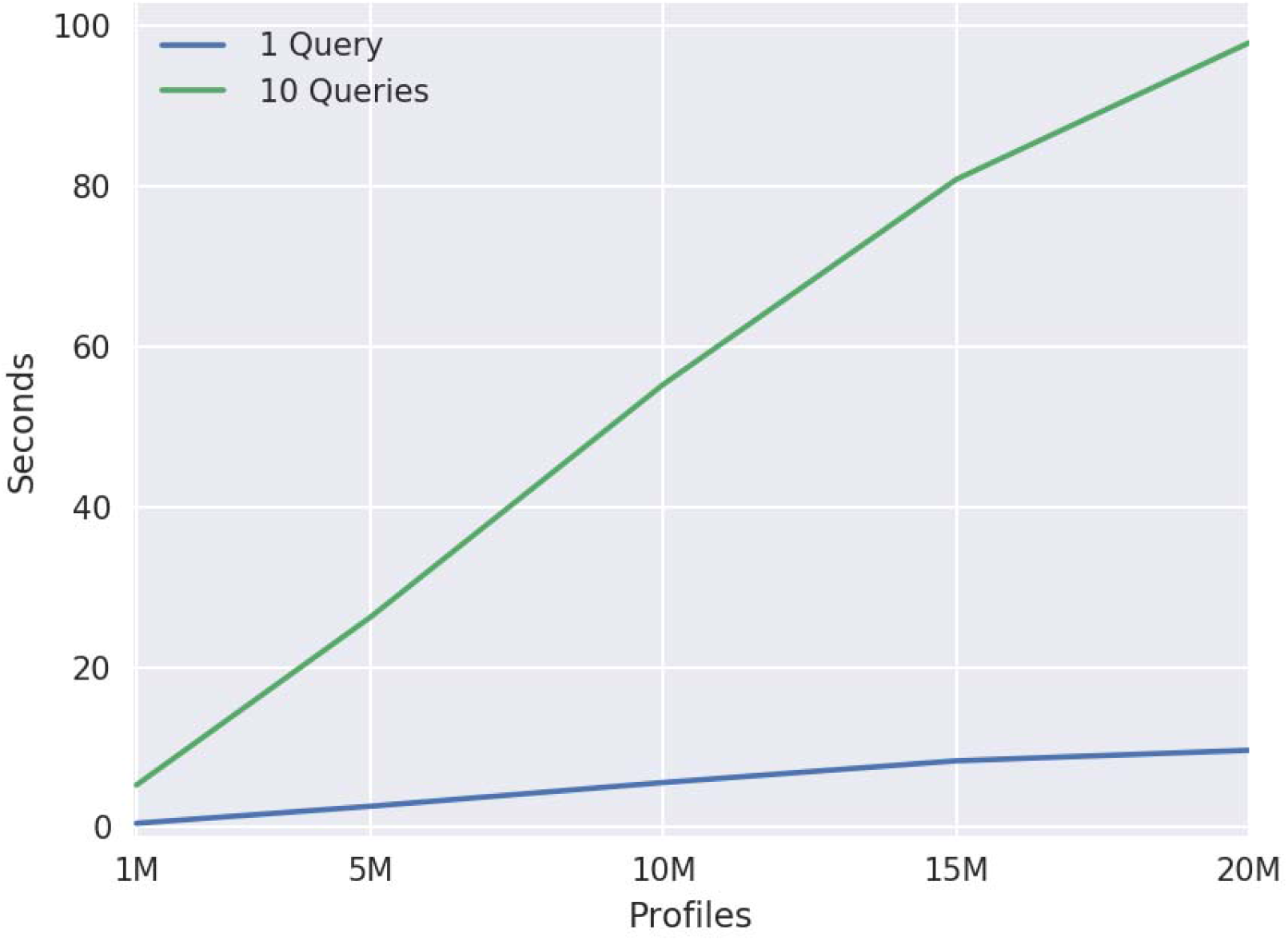
FastID profile comparison times for profile searches for 5,000 SNP panels. FastID was run on a SAGER laptop with 4.0 GHz i7 6700 CPU with a single computational thread.

SNPs with a targeted range of minor allele frequencies (MAF) for kinship analysis were selected from the panel of 39k SNPs from ALFRED. The number of mismatches between close relatives is expected to be lower than between unrelated individuals. The results for an individual compared to 4 generations of descendants and one million unrelated individuals are shown in Figure 2–4 using SNPs with MAF in the range of 0.01 and 0.2. The kernel density estimation (KDE) in statistics is a data smoothing calculation on a variable; this KDE graphs illustrates multiple peaks for both related and unrelated individuals (Figure 2). The source for differences between related individuals is coupled to the degree of relatedness (Figure 3) and impact of admixture (Figure 4) on comparison results. The peak distributions for the unrelated individuals reflect ethnicity differences with the left-most peak distribution reflecting unrelated individuals with the same ethnicity as the individual being searched with; the peaks further to the right reflect results comparing to other ethnicities. These results provide insights into the impact of ethnicity on kinship analyses, for relatives with the same degree of relatedness, relatives with shared ethnicities will appear closer than relatives of the same degree of relatedness but with ethnic differences.

**Figure 2.**
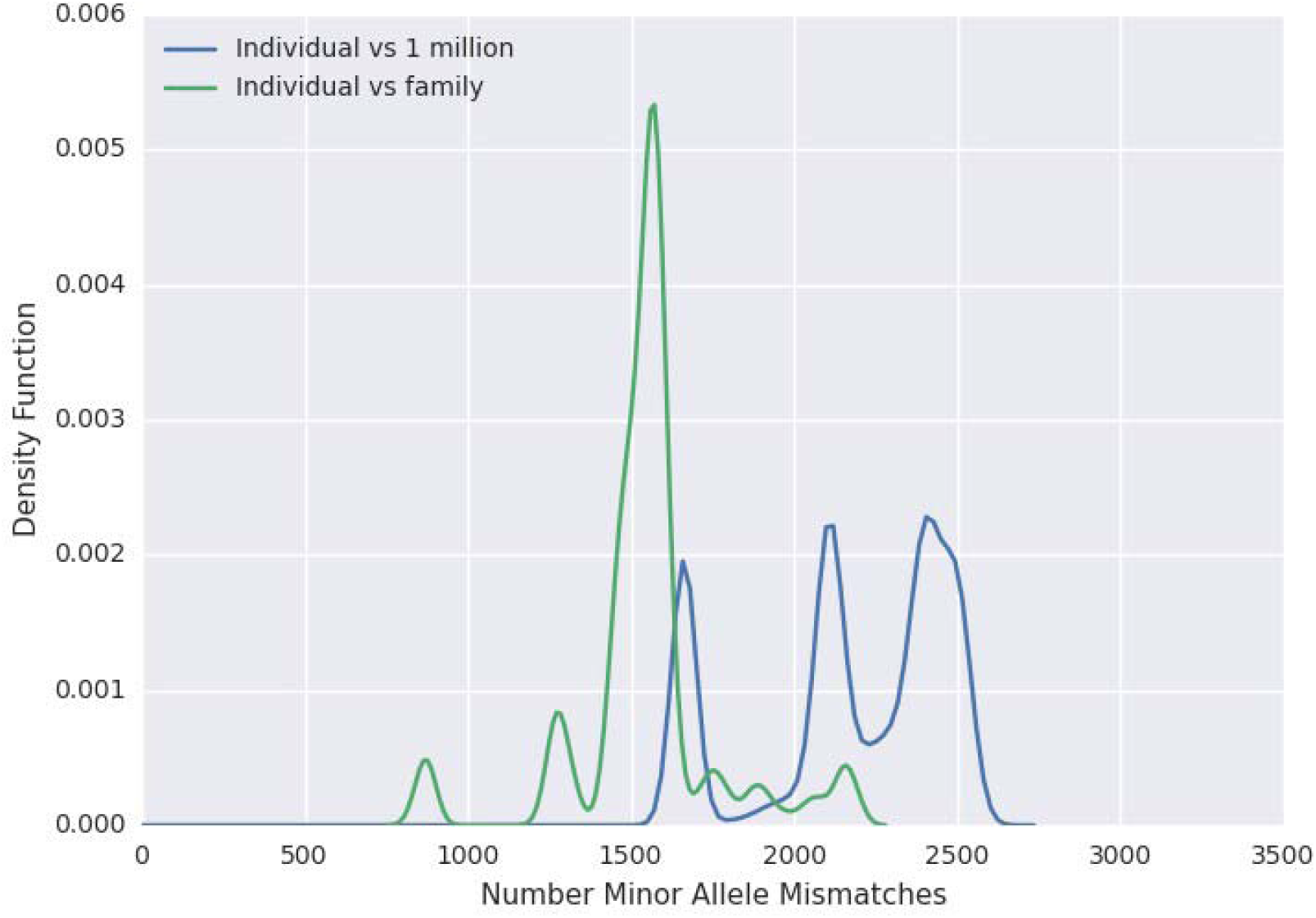
Comparison results for relatives (green) and unrelated individuals (blue) for an individual for 12,456 SNPs with a MAF between 0.01 and 0.2.

**Figure 3.**
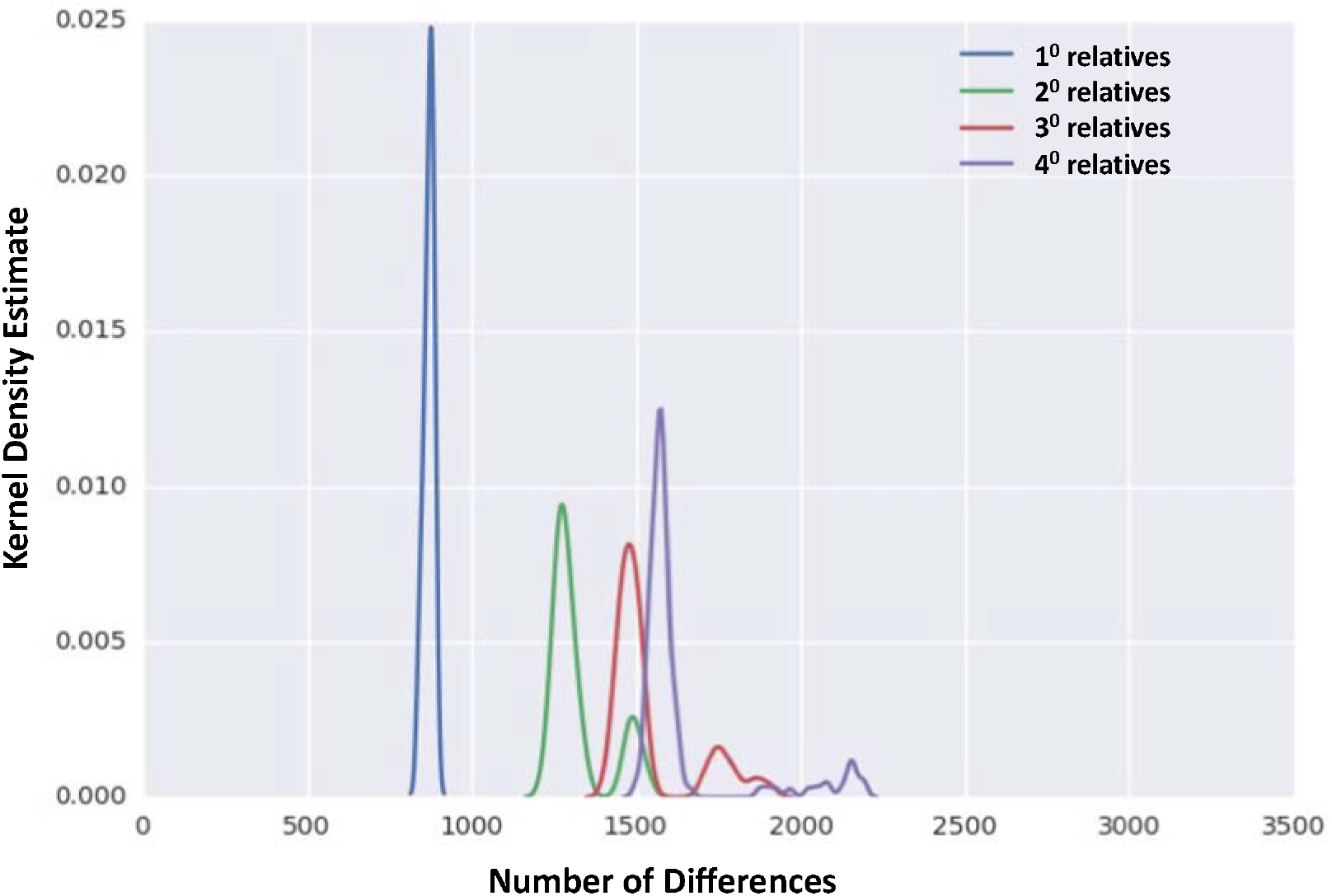
KDE comparison results for relatives for 12,456 SNPs with a MAF between 0.01 and 0.2. The first-degree relatives are children, the second-degree relatives are grandchildren, the third-degree relatives are great grandchildren, and the forth degree relatives are great great grandchildren.

**Figure 4.**
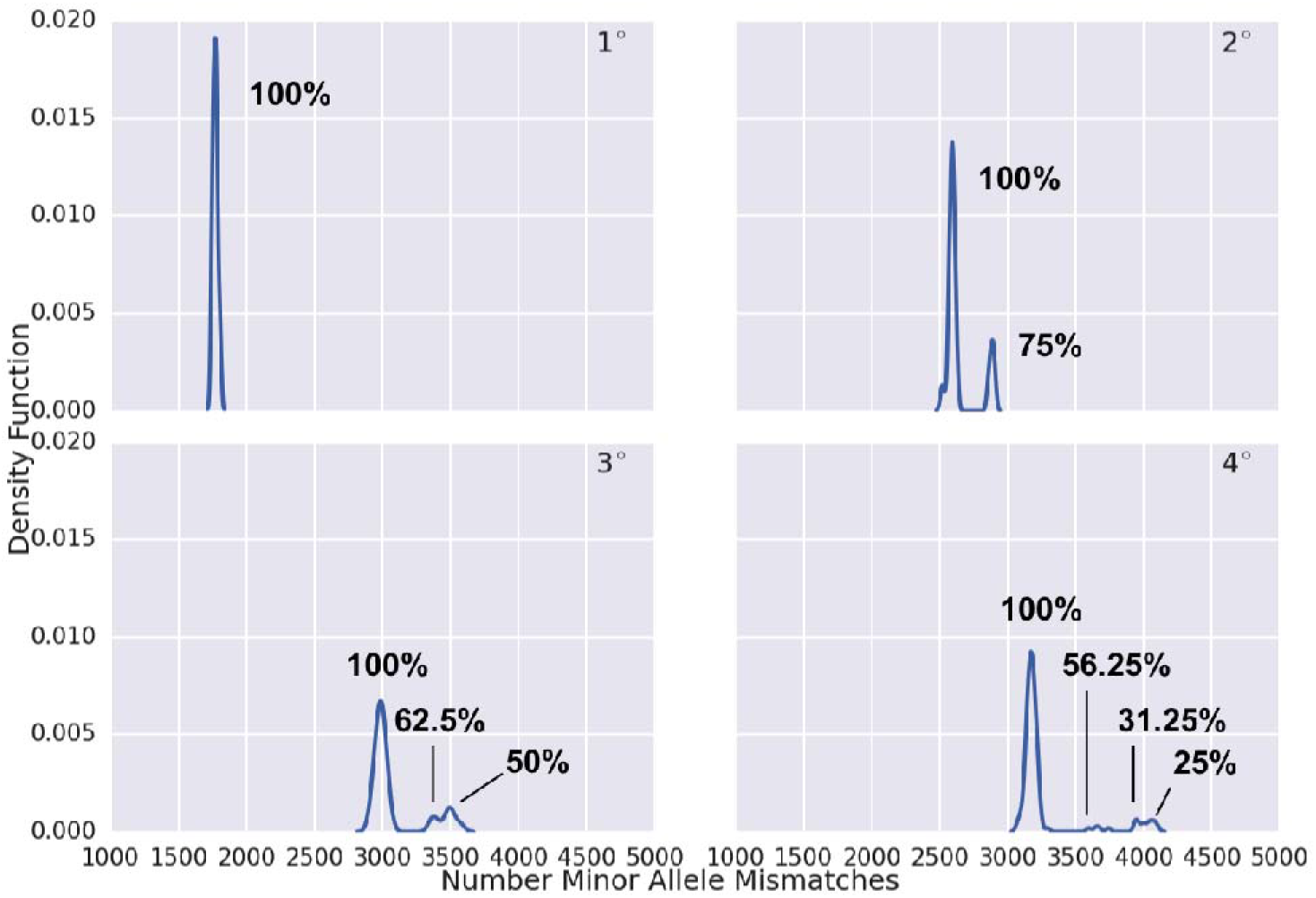
KDE results for close relatives including admixture. (Percentages identify the fraction of progenitor ethnicity shown in the distribution) for 12,456 SNPs with a MAF between 0.01 and 0.2.

## Summary

This in silico reference dataset uses SNP information from four different ethnic groups, with multiple generations that include intra-, and inter-ethnic marriages. The database tracks eleven million individuals with 39,108 SNPs, and familial relationships across eight generations. These datasets are available to aid development and testing of forensic software for identification, like FastID^15^, kinship modeling^16^, admixture, and likely additional applications.

## Data availability

The 11 million SNP profiles datset is available in Ricke, Darrell, 2018, “11 Million SNP Profiles datasets”, https://doi.org/10.7910/DVN/IDT8HZ, Harvard Dataverse.

## Footnotes

1 Distribution Statement A: Approved for public release: distribution unlimited. This material is based upon work supported under Air Force Contract No. FA8721–05-C-0002 and/or FA8702–15-D-0001. Any opinions, findings, conclusions or recommendations expressed in this material are those of the author(s) and do not necessarily reflect the views of the U.S. Air Force.

